# CAMP: a Convolutional Attention-based Neural Network for Multifaceted Peptide-protein Interaction Prediction

**DOI:** 10.1101/2020.11.16.384784

**Authors:** Yipin Lei, Shuya Li, Ziyi Liu, Fangping Wan, Tingzhong Tian, Shao Li, Dan Zhao, Jianyang Zeng

**Affiliations:** Machine Learning Department, Silexon AI Technology Co.Ltd., Nanjing, China; Institute for Interdisciplinary Information Sciences, Tsinghua University, Beijing, China; MOE Key Laboratory of Bioinformatics, TCM-X Center, Bioinformatics Division, BNRIST, Department of Automation, Tsinghua University, Beijing, China; MOE Key Laboratory of Bioinformatics, Tsinghua University, Beijing, China

**Author notes:** Corresponding authors: Dan Zhao, and Jianyang Zeng,.

**Keywords:** Peptide-protein interaction, peptide binding site identification, convolution neural network, self-attention

## Abstract

Peptide-protein interactions (PepPIs) are involved in various fundamental cellular functions and their identification is crucial for designing efficacious peptide therapeutics. To facilitate the peptide drug discovery process, a number of computational methods have been developed to predict peptide-protein interactions. However, most of the existing prediction approaches heavily depend on high-resolution structure data. Although several deep-learning-based frameworks have been proposed to predict compound-protein interactions or protein-protein interactions, few of them are particularly designed to specifically predict peptide-protein interactions. In this paper, We present a sequence-based **C**onvolutional **A**ttention-based neural network for **M**ultifaceted prediction of **P**eptide-protein interactions, called **CAMP**, including predicting binary peptide-protein interactions and corresponding binding residues in the peptides. We also construct a benchmark dataset containing high-quality peptide-protein interaction pairs with the corresponding peptide binding residues for model training and evaluation. CAMP incorporates convolution neural network architectures and attention mechanism to fully exploit informative sequence-based features, including secondary structures, physicochemical properties, intrinsic disorder features and position-specific scoring matrix of the protein. Systematical evaluation of our benchmark dataset demonstrates that CAMP outperforms the state-of-the-art baseline methods on binary peptide-protein interaction prediction. In addition, CAMP can successfully identify the binding residues involved non-covalent interactions for peptides. These results indicate that CAMP can serve as a useful tool in peptide-protein interaction prediction and peptide binding site identification, which can thus greatly facilitate the peptide drug discovery process. The source code of CAMP can be found in https://github.com/twopin/CAMP.

## 1 Introduction

Peptides play crucial roles in human physiology by interacting with a variety of proteins and participating in many cellular processes, such as programmed cell death, gene expression regulation and signal transduction [1, 2]. Owing to their safety, favorable tolerability profiles in human bodies and good balance between flexibility and conformational rigidity, peptides have become good starting points for the design of novel therapeutics, and identifying accurate peptide-protein interactions (PepPIs) is crucial for the invention of such therapeutics. Despite this fact, it is generally time-consuming and costly to determine peptide-protein interactions experimentally [1, 3]. To mitigate this issue, a number of computational methods have been developed to facilitate peptide drug discovery.

Sequence-based methods and structure-based methods are two mainstream approaches for protein-ligand interaction prediction. Sequenced-based methods mainly exploit primary sequence information to model the interactions. For example, CGKronRLS [4] and NRLMF [5] calculate sequence similarities and then use machine learning models to predict interactions between proteins and their ligands. These methods often require known protein-ligand interactions as supervised labels and pairwise similarity scores of proteins (or ligands) as input features, which is often impractical for large-scale data due to the huge computational complexity of similarity calculation. In addition, these approaches are not able to identify crucial binding residues, which hits a roadblock in deciphering the underlying mechanisms of PepPIs. Structure-based methods, such as molecular docking inherently tackle the problem by modeling structural poses at atom level and predicting binding affinities. There are many well-established docking strategies for determining PepPIs, which can be roughly divided into local (e.g., DynaRock [6] and Rosseta FlexPepDock [7]) and global docking methods (e.g., PIPER-FlexPepDock [8] and HPEPDOCK [9]) according to the extent of input structural information. Most of these docking approaches require three dimensional (3D) structure information to calculate binding free energies. Unfortunately, solving such 3D structures is generally time-consuming and expensive [1], letting alone consuming a large amount of computational resources due to the high computational complexity of the energy functions.

More recently, the booming deep learning technologies have provided feasible solutions to model protein-ligand or protein-protein interactions with better accuracy while requiring less computational resources. For instance, Cunningham et al. developed a hierarchical statistical mechanical modeling (HSM) approach [10] to predict the interactions between peptides and protein binding domains (PBDs). Wan et al. developed DeepCPI [11], a powerful computational framework that combines representation learning with a multi-modal neural network to predict compound-protein interactions (CPIs), and Chen et al. presented a siamese residual recurrent convolutional neural network (RCNN) [12] to predict protein-protein interactions.

Although the peptide drugs have increasingly attracted immense attention and the number of approved peptide therapeutics has been on the incline over the recent decades, only a few works have been proposed to exploit machine learning or deep learning methods to model peptide-protein interactions. Furthermore, for deciphering the underlying mechanisms of peptide-protein interactions, the existing approaches mainly focus on identifying peptide-binding residues on protein surface, such as the sequence-based method PepBind [13] and the structure-based method InterPep [14]. PepBind [13] is a sequence-based method for peptide-binding residue prediction, which assumes that a protein would have fixed binding residues even interacting with different peptides. However, in many cellular processes, different peptides with diverse biological functions may present distinct binding poses to a single protein, which thus may involve different protein residues in the interaction. Therefore, PepBind intrinsically fails to model the situations that multiple peptides interacted with different regions of a protein surface [13]. InterPep combines a random forest model with hierarchical clustering to predict the regions of a protein structure where the input peptide is most likely to bind [14], which requires a target protein structure and a peptide sequence, and thus its application may be limited to only those proteins with available 3D structural data.

Moreover, most of existing computational methods in modeling peptide-protein interactions fail to answer an important question, which is frequently raised by pharmacologists — how to determine the contribution of each individual peptide residue to the binding activity? Therefore, there is a manifest need for addressing the following challenges: (1) identifying the peptide-protein interactions accurately and efficiently, taking account of information from both peptides and proteins; (2) possessing the great generalization ability to large datasets; and (3) detecting crucial binding sites of peptides that can provide useful hints for downstream amino acid substitution or backbone modification.

In this paper, we introduce CAMP, a sequence-based deep learning framework for both peptide-protein interaction prediction and peptide binding residue identification. CAMP first exploits a series of sequence-based features, including secondary structures, physicochemical properties, intrinsic disorder scores and the position-specific scoring matrix of the protein. These predefined features generally summarize a group of informative biophysical properties of primary sequences of proteins and peptides [15–20]. Next, our deep learning architecture processes the encoded features through embedding modules containing self-learnt word embedding layers [21] and dense modules containing fully-connected layers to capture latent features, which are then fed into convolution neural network (CNN) and attention modules to leverage both sequential and local information. Finally, the outputs of these modules are combined together to predict peptide-protein interactions. In addition, CAMP generates a predicted score for each peptide residue, suggesting whether this residue is a binding site. This prediction can not only provide useful insights for characterizing essential residues involved binding activities, but also enhance the binary interaction prediction by providing extra supervised information.

In summary, the major contributions of this work lie in the following three perspectives:

1. We develop a sequence-based deep learning framework for both peptide-protein interaction prediction and peptide binding site identification. From the input side, features are generated based on the primary sequences of peptides and proteins, which thus can relieve the dependence of 3D structure data. To our best knowledge, our work is the *first* sequence-based deep learning framework to predict peptide-protein interactions as well as identify peptide binding sites involved in the interactions.
2. We evaluate the predicting power and generalization ability of CAMP on a benchmark dataset and also an independent test dataset from the RCSB Protein Data Bank (PDB) [22, 23] and DrugBank [24–28] for both binary peptide-protein interaction prediction and peptide binding site identification.
3. We extend the applications of CAMP by further exploring its architecture to address a similar task — peptide-PBD (protein binding domain) interaction prediction. We trained CAMP using peptide-PBD interaction data and performed cross-validation for eight PBD families. CAMP outperformed HSM, a bespoke machine learning framework for peptide-PBD interaction prediction, which thus verifies the robustness and the wide application potential of CAMP.

## 2 Results

### 2.1 CAMP exploits sequence-based features for multifaceted PepPI prediction

CAMP provides a sequence-based deep learning framework for simultaneously predicting peptide-protein interactions and peptide binding residues given an input pair of peptide and protein sequences. CAMP first generates a group of sequence-based features as input (Section 4.3, Supplementary Notes S1 and Figure S1), which are calculated from the primary sequences of peptides and proteins. Next, CAMP predicts binary interaction and peptide binding residues for a given peptide-protein pair through a convolutional attention-based neural network (Section 4.4). As shown in Figure 1, CAMP has six groups of modules: (1) embedding modules containing self-learnt word embedding layers [21] that take protein and peptide sequence information as input separately with categorical features, including amino acid, secondary structures (SS-based) and polarity and hydropathy property (physicochemical) representations of peptides and proteins; (2) dense modules containing fully-connected layers that take numerical features, including intrinsic disorder (ID-based) representations of peptide and protein sequences and PSSM representations of protein sequences; (3) convolution neural network (CNN) modules that extract local contextual features and global sequence information for both peptides and proteins; (4) self-attention modules that learn the contributions of individual residues from protein and peptide sequences to the final prediction; (5) a peptide binding site prediction module that takes the output of the peptide convolutional layers with a sigmoid activation function for each position to predict whether each peptide residue binds to the partner protein; and (6) a binary interaction prediction module that combines all the extracted features of peptides and proteins and uses three fully-connected layers to finally predict whether there exists an interaction between a given peptide-protein pair.

**Fig. 1.**
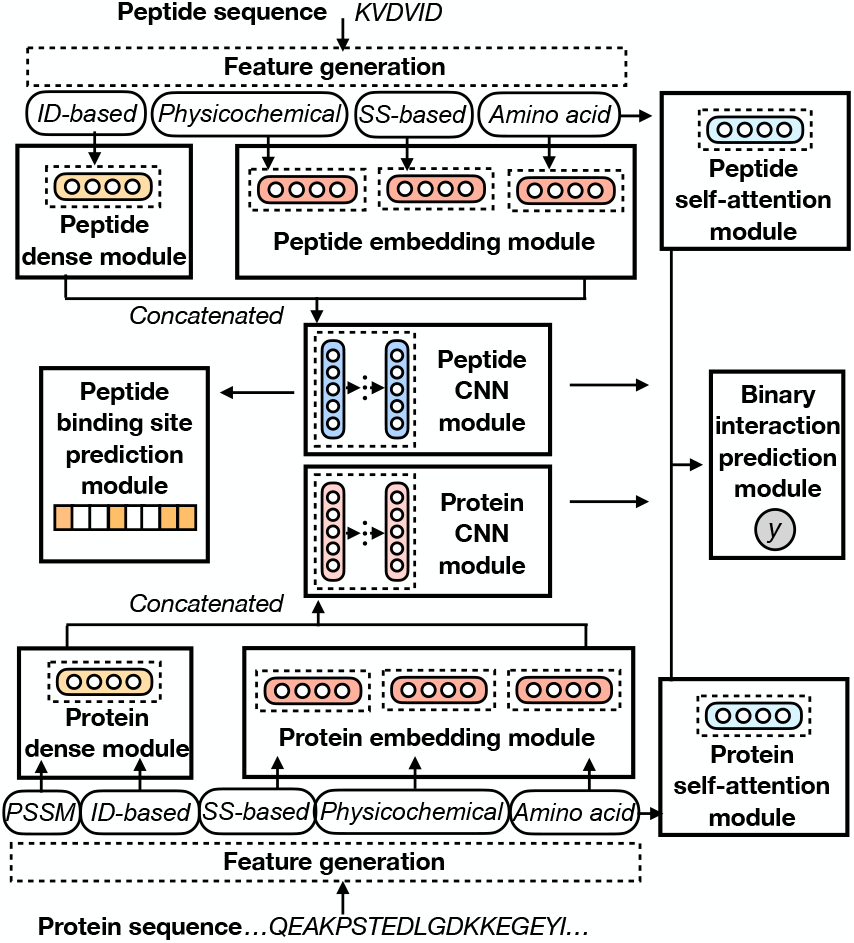
The network architecture of CAMP. Given a peptide-protein pair, we first generate sequence-based features (Section 4.3). For the input peptide sequence, its amino acid, physicochemical and secondary structure-based (SS-based) representations are separately fed into three embedding layers and its intrinsic disorder-based (ID-based) representation is fed into a dense layer for feature extraction. Then these features are concatenated together and fed into a CNN module. The features of the input protein sequence including its amino acid, physicochemical and SS-based representations, are also fed into three embedding layers and the dense layer takes the ID-based and PSSM representations as inputs. Next, the outputs of these modules are concatenated together and fed into CNN modules and the output of the amino acid representations of the peptide and the protein are fed into two self-attention modules to learn the importance of individual residues (i.e., the contributions of individual residues to the final prediction). After that, the outputs of self-attention modules and CNN modules are concatenated together to predict a binding score for each peptide-protein pair through three fully-connected layers and a binding score for each residue from the peptide sequence using the outputs of the peptide CNN module.

### 2.2 CAMP outperforms baseline methods in pairwise binary interaction prediction

The binary classification of peptide-protein interactions is the primary goal of CAMP. Here we compared the classification performance of CAMP with that of other state-of-the-art baseline methods, including a similarity-based matrix factorization method called NRLMF [5], a deep-learning-based model for protein-protein interaction (PPI) prediction called PIPR [12], and a deep-learning-based model for compound-protein interaction (CPI) prediction called DeepDTA [29]. All the prediction methods were evaluated on a benchmark dataset through cross-validation (Section 4.1). The area under the receiver operating characteristics curve (AUC) and the area under precision recall curve (AUPR) were used to evaluate the performance of all models. In general, AUPR can provide a better metric to evaluate the prediction models on skewed data in a more informative way than AUC [30]. Since the human-curated data may contain “redundant” interaction pairs (e.g., one protein interacting with more than one similar peptide or vice versa), which could be easily predicted by the models. To avoid the trivial predictions caused by such cases, we followed the same strategy as in MONN [31], and mainly used the cluster-based cross-validation settings for performance evaluation. In particular, based on similarity scores derived from Smith-Waterman alignment (https://github.com/mengyao/Complete-Striped-Smith-Waterman-Library), we divided proteins and peptides into different clusters such that the entities from the same cluster did not appear in the training and testing sets at the same time (Supplementary Notes S2). We evaluated the performance of CAMP and the baseline methods under three cluster-based cross-validation settings. More specifically, in the “new protein setting”, no proteins from the same cluster appeared in both training and testing sets; in the “new peptide setting”, no peptides from the same cluster appeared in both training and testing sets; and in the “both new setting”, neither proteins nor peptides from the same cluster appeared in training and testing sets at the same time. Figure 2 shows that CAMP consistently outperformed the state-of-the-art baseline methods, with an increase by up to 10% and 15% in terms of AUC and AUPR, respectively. In addition, we observed a slight decreasing trend of prediction performance for all methods with larger clustering thresholds, which generally corresponded to more difficult tasks. We also noticed that the model performance under the “new peptide setting” seemed to be better than that in the other settings. This can be explained by the fact that the peptides in our benchmark set shared less similarity with each other than proteins, and thus the distributions of peptides in the training and testing sets did not change much after clustering based on similarities. Such test results suggested that CAMP can achieve better and more robust performance than the baseline methods under all cross-validation settings. We also conducted comprehensive ablation studies to demonstrate the importance of individual components in the model architecture (results can be found in Supplementary Notes S3, Table S6).

**Fig. 2.**
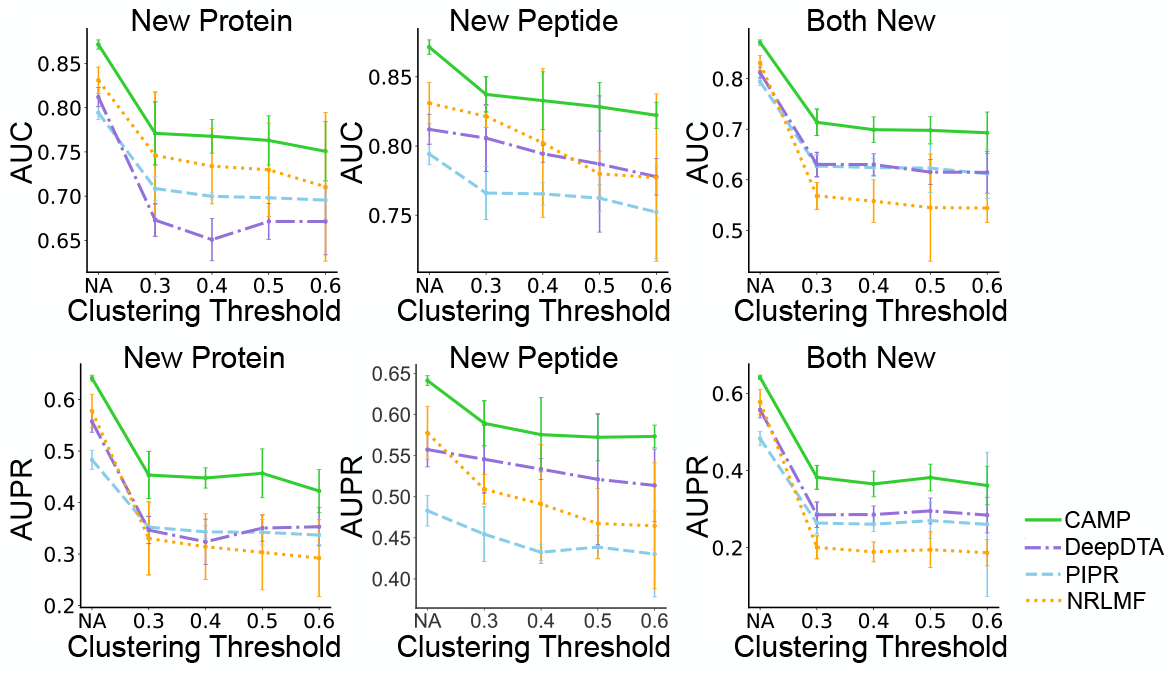
AUC and AUPR of CAMP and baseline models through five-fold cross-validation under “new protein setting” and “new peptide setting”, or nine-fold cross-validation under “both new setting” (definitions of different settings are described in Section 2.2). The error bars represent the standard deviations over all folds. “NA” stands for random cross-validation, i.e., randomly splitting the dataset and used 80% of the dataset to train the model and the remaining 20% to evaluate the performance.

### 2.3 CAMP provides new insights about peptide-protein interactions by identifying binding residues on peptides

So far a number of computational methods have been developed for predicting the binding sites on the protein surface in PepPI predictions [32, 14, 33]. These methods learn from 3D structure information of peptide-protein complexes and can pinpoint binding sites on protein surface with relatively good accuracy. However, few models are specifically designed to characterize binding sites on the peptides in PepPIs, which are also crucial for understanding the biological roles of peptides and designing efficacious peptide drugs. For pharmacologists, the choice of chemical modification heavily relies on the identification of essential peptide residues involved in binding activities [1]. Conventionally, pharmacologists would iteratively replace possible residues and conducted wet experiments for verification. Although these attempts could provide useful information for further drug design, e.g., changing particular non-binding residues or modifying groups on their side chains to improve stability and reduce toxicity [2, 1], these experimental approaches are generally expensive and time-consuming.

In CAMP, we designed a supervised prediction module to identify binding residues from a peptide sequence. We first constructed a set of qualified labels for peptide binding residues using the interacting information derived from PepBDB [34], which is a comprehensive structure database containing the known interacting peptide-protein complexes from the RCSB PDB [22, 23, 35] and information about interacting residues in peptides involved in hydrogen bonds and hydrophobic contacts. With the support from such supervised information, CAMP achieved an average AUC of 0.806 and Matthews Correlation Coefficient (MCC) (definitions can be found in Supplementary Notes S4) of 0.514 on peptide binding site prediction using a five-fold cross-validation procedure under the “random-split setting” (Figure 3A and 3B). We also illustrated that CAMP was able to yield relatively robust performance with respect to the peptide length and the numbers of binding residues on the peptide (detailed results can be found in Supplementary Figure S2).

**Fig. 3.**
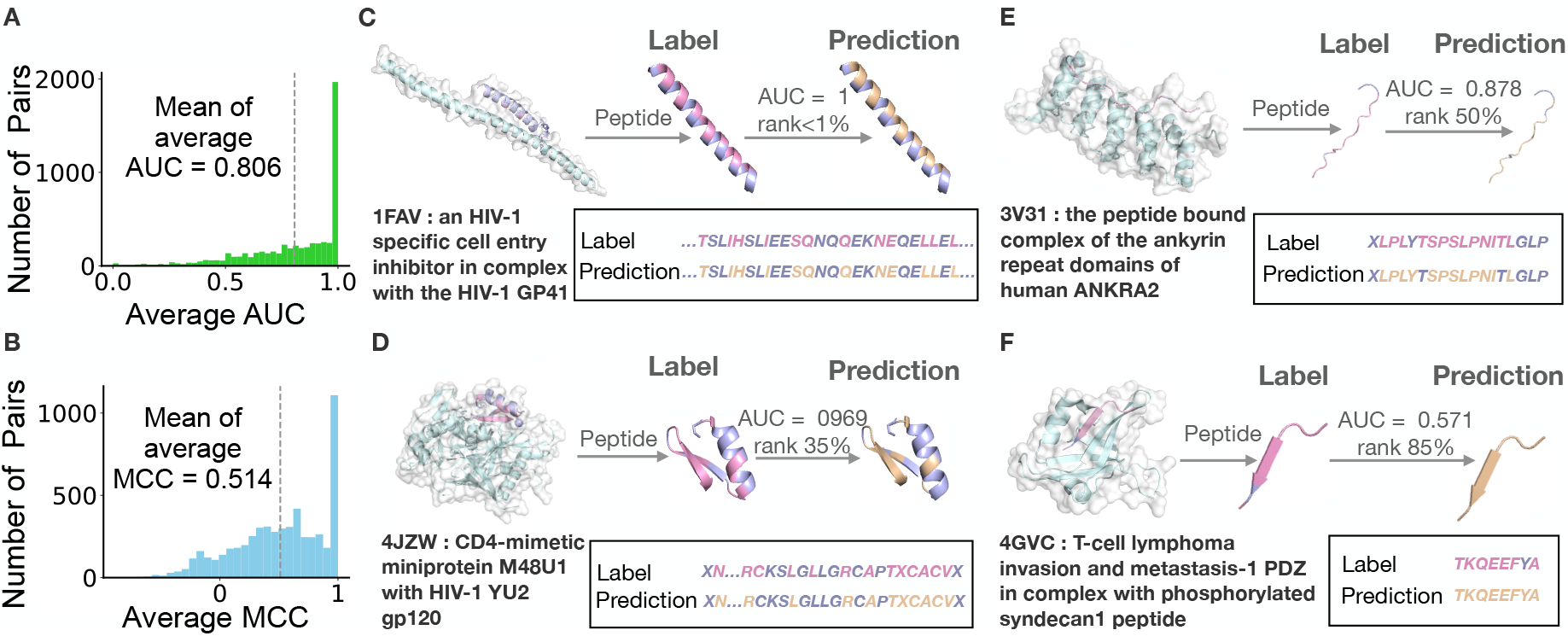
Performance evaluation of CAMP on peptide binding site prediction on the benchmark dataset through five-fold cross validation. (A) and (B) show the distributions of AUC and MCC for peptide binding site prediction, respectively. The mean values of average AUC and MCC are plotted in dotted lines. (C) - (E) show four examples of peptide binding site predictions by CAMP that ranked around 1%, 35%, 50% and 85% in terms of average AUC, respectively. The PBD complexes were retrieved from the RCSB PDB [22, 23, 35] and the images were generated by PyMOL [36]. The protein chains in the complexes are colored in lightblue while the peptide chains are colored in light purple and pink. For each peptide, the true binding residues are colored in pink while the predicted binding residues generated by CAMP are colored in wheat.

To further demonstrate the performance of CAMP in binding site prediction, we also selected four representative cases (ranked around 1%, 35%, 50% and 85% in terms of the average AUC scores of predicted peptide binding sites, respectively) and compared the predicted sites with the true binding ones. Figure 3C shows the first example, a complex of an HIV-1 specific cell entry inhibitor and HIV-1 GP41 trimeric core (PDB ID: 1FAV). The peptide inhibitor has 33 amino acids and 12 of them are binding residues. CAMP identified all these binding sites without any false positives. Such a prediction was the most ideal case in our prediction task and we found that 30.2% of the binding site predictions were completely accurate like this case. Figure 3D shows the second example, a complex of HIV-1 gp120 envelope glycoprotein and the CD4 receptor (PDB ID: 4JZW), which ranked around top 35% in terms of the average AUC. The peptide has 28 amino acids and 13 of them are binding residues. Our predicted binding residues covered 11 true binding residues along the peptide sequence and missed two true binding residues. Figure 3E shows the third example, a complex of a peptide from histone deacetylase and the ankyrin repeat family A protein (PDB ID: 3V31). This pair ranked around the median among our predictions in terms of AUC and 11/13 of the true binding residues were successfully identified by CAMP with one false positive. Figure 3F shows the last example, a complex of the T-lymphoma invasion and metastasis inducing protein and a 8-residue phosphorylated syndecan-1 peptide (PDB ID: 4GVC), which ranked around 85% among our predictions with an average AUC of 0.571. All eight residues including one false positive were predicted as binding residues by CAMP. Overall, our test results demonstrated that CAMP yields accurate binding site predictions and thus can provide reliable evidence for further understanding the interacting mechanisms of peptides with their partner proteins.

### 2.4 CAMP correctly identifies the GLP-1 receptor as a target of Semaglutide and its analogs

Glucagon-like peptide receptor (GLP-1R) agonists play an important role in the treatment of type 2 diabetes mellitus (T2DM) [37, 38]. We next investigated whether CAMP was able to correctly identify the interactions of Semaglutide, a known GLP-1R agonist (GLP-1RA), and its analogs with GLP-1R. In our benchmark data set, there are seven Semaglutide-analogous peptides that bind to GLP-1R. To avoid ‘easy prediction’, we removed those GLP1RA peptide drugs from the training set that shared similar sequences (defined as peptide sequence similarities > 40%) with Semaglutide (e.g., Liraglutide and Taspoglutide), and had interacting proteins similar to GLP-1R (i.e., with protein sequence similarities > 40%). After removing these records as well as seven pairs of Semaglutide-analogous peptides and GLP-1R, we re-trained the CAMP model and combined the seven Semaglutide-analogous peptides with the remaining 3,400 proteins to construct an independent test set which contained 23,800 candidate pairs. The test showed that CAMP was able to identify six of seven interacting pairs of Semaglutide-analogs peptides and GLP-1R with an AUC score of 0.831. For all the Semaglutide-analogs peptides, GLP-1R was ranked to the top 10% almost among all the candidate proteins (detailed results can be found in Supplementary Table S7). Such results further demonstrated the strong predictive power of CAMP.

We also examined the predicted binding residues of Semaglutide with its receptor (detailed results can be found in Supplementary Fig S3). CAMP correctly identified 11/12 of the true binding residues of Semaglutide with an average AUC of 0.917. Such a prediction result can provide useful insights for pharmacologists if they aim to improve the stability of the peptide drugs by replacing the non-binding residues with synthetic amino acids without changing the interacting interface of the binding complexes.

### 2.5 CAMP achieves great generalizability on an independent test dataset

We conducted an additional test to further illustrate the generalizability of CAMP on binary interaction prediction and peptide binding residue identification. In particular, we evaluated CAMP on an independent dataset derived from the PDB [39, 23, 35] following the same strategy as in constructing our previous benchmark dataset. This additional test set contained 379 PepPIs from 262 peptides and 246 proteins from the PDB complexes released from October 1st, 2019 to March 10th, 2020. We also randomly paired these peptides and proteins without known evidence of interactions in the test set to obtain negative samples.

To demonstrate the robust performance of CAMP on binary interaction prediction, we evaluated the performances CAMP and the baseline models on several variations of the test dataset with different positive-negative ratios. Each model was first trained on the complete benchmark dataset and then an ensemble version (i.e., average predictions from five models) was used to make predictions on the additional test datasets. Figure 4A and 4B show that CAMP achieved the best results under all scenarios, demonstrating that CAMP outperformed the baseline methods with a relatively robust performance. We also observed that the AUC of all methods increased slightly as the positive-negative ratio decreased from 1:1 to 1:10. This was probably because the increased sample size brought more information for models to learn. Also, the AUPR of all methods decreased more dramatically than AUC as the positive vs. negative ratio increased. This was mainly because AUPR is generally more affected by the ratio of positive vs. negative samples [30].

**Fig. 4.**
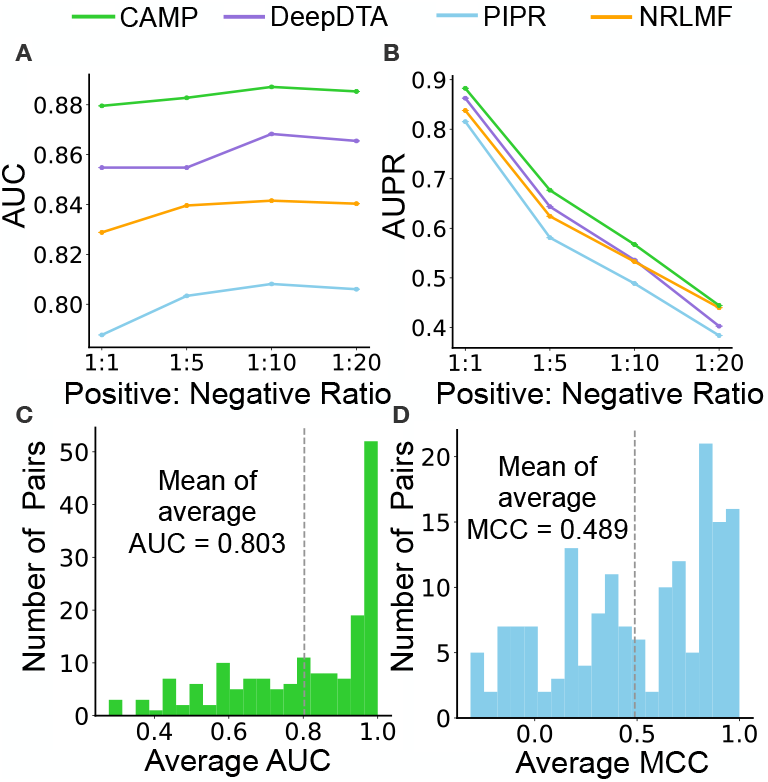
CAMP yielded robust performance and outperformed the baseline models on an independent test set. (A) and (B) show the evaluation results with different positive-negative ratios of the test dataset in terms of AUC and AUPR, respectively. (C) and (D) show the distributions of AUC and MCC for peptide binding site prediction, respectively. The mean values of average AUC and MCC are plotted with dotted lines.

We also evaluated the prediction results of CAMP on the identification of peptide binding residues. We obtained the annotated binding residues of peptide sequences from PepBDB [34]. In total, 208 PepPIs have such peptide binding residue labels from the test dataset. Figure 4C and 4D show that CAMP was able to maintain its prediction power on the additional dataset.

### 2.6 Performance of CAMP on predicting peptide-PBD interactions

Although we rarely found deep-learning-based methods for predicting PepPIs, there was a machine learning approach, called hierarchical statistical mechanical modeling (HSM) [10], focusing on a quite related problem, i.e., predicting the interactions between peptides and globular protein-binding domains (PBDs). The PBD-containing proteins play essential roles in a variety of cell activities, e.g., multiprotein scaffold formation and enzyme activity regulation [40–42]. By incorporating biophysical knowledge as prior information into a machine learning framework, HSM was reported to yield superior prediction performance on eight common PBD families with AUC scores ranging from 0.88 to 0.92. We compared CAMP with two reported models of HSM, i.e., HSM-ID (in which eight separate models were trained for each PBD/enzyme family) and HSM-D (in which a single unified model was trained for all families), on predicting peptide-PBD interactions.

Here, we compared the performance of CAMP with that of HSM models on predicting peptide-PBD interactions. In particular, we evaluated the performance of CAMP with the same dataset and eight-fold cross-validation setting as used in the HSM paper (see Supplementary Notes S5 for more details). Figure 5 shows that CAMP significantly outperformed both HSM-ID and HSM-D across all domain families except the PDZ family. We also noticed that HSM-ID and HSM-D had large prediction variation across different families. As explained in the HSM paper, this may due to the skewed distribution of the data (i.e., the numbers of pairs from different families were imbalanced). For families of large data amount like PDZ, the HSM models could learn quite well but for those families of relatively small data sizes like domains from the phosphotyrosine binding (PTB) family, HSM models had obvious drop in performance. In contrast, the performance of CAMP was more robust and less influenced by the fluctuant data sizes. Such results indicated that CAMP is also suitable for tackling the related peptide-PBD interaction prediction problem.

**Fig. 5.**
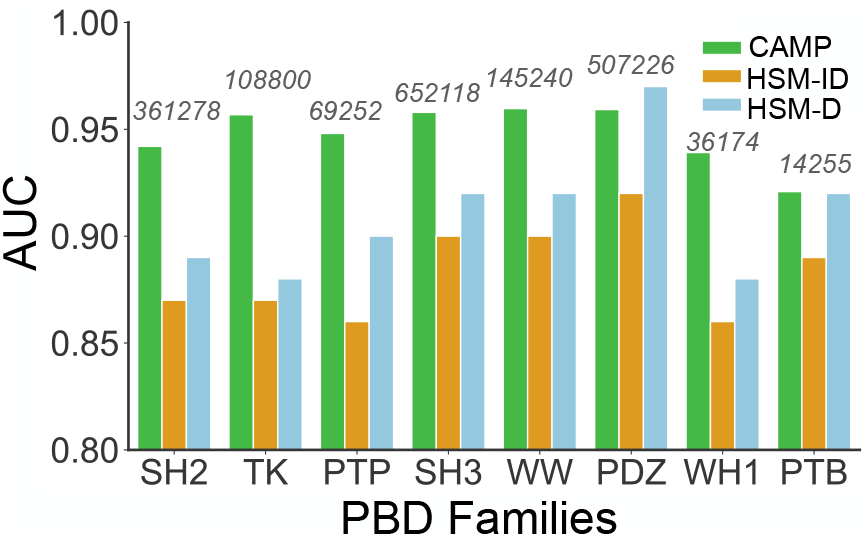
Model performance of CAMP, HSM-ID and HSM-D across eight families. CAMP achieved a relatively stable performance over all families while the performances of HSM models were easily influenced by the sample size (marked in grey number) in the training set. CAMP outperformed the HSM models, with an increase of AUC by 3% – 7%. All the evaluation metrics of the HSM models were obtained from the origin paper [10].

## 3 Conclusion

In this work, we proposed a sequence-based deep learning approach CAMP for multifaceted peptide-protein interaction prediction, including binary interactions prediction as well as peptide binding residue identification. We first generated a series of sequence-based features for both peptides and proteins. Compared to traditional peptide or protein feature encodings such as k-mer representation, our sequence-based representations combined informative structurally-annotated features, protein evolutionary profiles and intrinsic disorder scores of the peptide to enhance the peptide-protein interaction prediction. Comprehensive cross-validation evaluation demonstrated the superior performance of CAMP over the state-of-the-art baseline methods on binary interaction prediction. Furthermore, we seeked to decipher the underlying mechanisms of peptide-protein interactions by identifying the peptide binding residues. We showed that CAMP can accurately detect the binding residues from the peptide sequence. We also presented four representative cases to visualize the results of peptide binding site prediction and examined the predicted targets for Semaglutide and its analogs. In addition, we further verified the prediction power of CAMP by extending its applications to a peptide-protein binding domain (PBD) interaction prediction task. All these results indicated that CAMP can provide accurate peptide-protein interaction predictions as well as useful insights into understanding the peptide binding mechanisms.

Nevertheless, there still exist certain limitations in the current version of CAMP. For example, it cannot directly predict the binding residues from the protein sequence in a given peptide-protein pair. Proteins usually have much longer sequence lengths than peptides, which will thus introduce more parameters for CAMP to learn. More specifically, in our problem setting, peptides only have 50 amino acids at most while protein sequences may have more than 1,000 amino acids. It would be much more challenging for CAMP to capture the binding residues among hundreds of protein residues. In addition, the labels of protein binding residues can only be retrieved based on the PDB complexes, which often contain fragments of interacting proteins instead of unified intact sequences. The gap between the protein fragments from PDB complexes and the unified protein sequences from UniProt [43] may hinder the prediction of protein binding residues.

## 4 Methods

### 4.1 Datasets

We constructed a benchmark dataset from two sources, i.e., protein-peptide complex structures from the RCSB Protein Data Bank (PDB) [22, 23, 35] and the known drug-target pairs from DrugBank [24–28] (more details of data curation can be found in Supplementary Notes S6). In total, we obtained 7,417 positive interacting pairs covering 3,412 protein sequences and 5,399 peptide sequences. Among them, 6,581 pairs from the RCSB PDB have residue-level binding labels in peptide sequences. We then constructed a negative dataset by randomly shuffling those non-interacting pairs of proteins and peptides. More specifically, for each positive interaction, five negatives were generated by randomly sampling from all the shuffled pairs of non-interacting proteins and peptides. Overall, we obtained 44,502 peptide-protein pairs as our benchmark dataset.

### 4.2 Problem formulation

In our problem setting, we mainly consider those peptide sequences with no more than 50 amino acids and protein sequences with more than 50 amino acids. We use 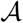 to denote a vocabulary of 21 types of amino acids (i.e., 20 canonical amino acids and a letter ‘X’ for any unknown or non-standard amino acid). Then, a given peptide-protein pair (*S*_*pep*_, *S*_*pro*_) can be defined as two sequences of amino acids *S*_*pep*_ = (*p*_1_, *p*_2_,…, *p*_*m*_), *S*_*pro*_ = (*q*_1_, *q*_2_,…, *q*_*n*_), in which each 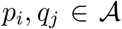 stand for the residue at position *i* of the peptide and position *j* of the protein, respectively, and *m, n* represent the lengths of the peptide and protein sequences, respectively.

Our sequence-based neural network model, CAMP, addresses two prediction tasks: (1) a binary classification task to predict PepPIs; (2) a binding site classification task to identify binding sites from the input peptide sequence. More specifically, the first prediction task can be described as a binary classification problem, in which label *y*_*i*_ = 1 indicates the existence of an interaction between the *i*-th peptide-protein pair and *y*_*i*_ = 0 otherwise. The output probability of CAMP for this task can be denoted by a real value between 0 and 1. The second prediction task aims to pinpoint the binding residues from the peptide sequence in a given peptide-protein pair. Here, for a peptide with *m* residues, we define its binding vector as ***b***_***pep***_ = (*b*_1_, *b*_2_,…, *b*_*m*_), in which each binary element *b*_*i*_ denotes whether the *i*-th residue binds to the partner protein (1 for the existence of binding and 0 otherwise).

### 4.3 Feature encoding

Our framework only requires sequence information as input, therefore alleviating the problem of limited structure data. Below we will describe how we encode the features of protein and peptide sequences.

#### Amino acid representations of peptides and proteins

The most common representations of proteins are amino acids. Here we define an alphabet of 21 elements to describe different types of amino acids (i.e., 20 canonical amino acids and a letter ‘X’ for unknown or non-standard ones). Each type of amino acid is encoded with an integer between 1 and 21. For each amino acid sequence *S* = (*a*_1_, *a*_2_,…, *a*_*n*_), we generate an *n* × 1 array, in which in the corresponding residue position, each element is an integer representing the amino acid type.

#### Secondary structure-based representations of peptides and proteins

Although our problem setting assumes that 3D structure data is unavailable, previous studies have suggested that the predicted structures of the amino acid sequences could still provide useful information [44, 16, 45]. Here, for each amino acid sequence *S* = (*a*_1_, *a*_2_,…, *a*_*n*_), we use SSPro [16] to generate an *n* × 1 array, in which each element is an integer representing the combination of secondary structure class and amino acid type at the corresponding position (see Supplementary Notes S1).

#### Physicochemical property representations of peptides and proteins

The hydrophobicity, hydrophilicity and polarity of the R groups of individual amino acids can affect the tendency of the interactions between residues [46]. For each amino acid sequence *S* = (*a*_1_, *a*_2_,…, *a*_*n*_), we generate an *n* × 1 array, in which each element is an integer representing the combination of the polarity and hydropathy properties of the residue at the corresponding position (see Supplementary Notes S1).

#### PSSM representations of proteins

Position-specific scoring matrices (PSSMs) are popular representations of protein sequences, which can detect remote homology of the protein sequences [47, 20]. For each protein sequence *S* = (*a*_1_, *a*_2_,…, *a*_*n*_) of length *n*, we use PSI-BLAST [19] to generate a normalized position-specific scoring matrix, an *n* × 20 array ***S***, in which each element *S*_*i,j*_ stands for the probability of the *j*-th amino acid type at position *i* in the protein sequence (see Supplementary Notes S1).

#### Intrinsic disorder-based representation of peptides and proteins

It has been reported that the intrinsic disorder-based features in peptide and protein sequences play an crucial role in protein-peptide interactions [15]. Here, for individual residues in the peptide and protein sequences, we first employ IUpred2A [18, 17] to predict its intrinsic disorder properties. For an amino acid sequence *S* of length *m*, we construct an *m* × 3 array representing three types of disorder scores for individual residues (see Supplementary Notes S1).

### 4.4 Network architecture

As mentioned in Section 2.1, our model consists of six groups of modules: (1) embedding modules that encode categorical features of protein and peptide sequences; (2) dense modules that encode numerical features of the peptide and the protein sequences; (3) convolution neural network (CNN) modules that extract latent information from peptide and protein features separately; (4) self-attention modules that learn the importance of each residue from protein and peptide sequences; (5) a peptide binding site prediction module to infer whether each peptide residue binds to the partner protein; and (6) a binary interaction prediction module to finally predict whether there exists an interaction between a given peptide-protein pair.

#### Embedding module and dense module

CAMP consists of two embedding modules as feature encoders, which take protein and peptide features as input separately (Figure 1). Each embedding module consists of three self-learnt word embedding layers [21], taking amino acid, secondary structure and physiochemical representations as input, respectively. In addition, each dense module consists of a fully-connected layer to take numerical features as inputs, i.e., the intrinsic disorder-based features (ranging between 0 and 1) of peptides and proteins as well as the normalized profile matrices (PSSM) of proteins. The outputs of the embedding layers and the dense layers are then concatenated together for both peptides and proteins.

#### The convolution neural network module

We deploy a popular deep learning architecture, convolution neural network (CNN), to extract the informative knowledge from the input sequence-based features. The CNN architecture is able to integrate local dependencies to capture latent information of sequential features and has been successfully used to predict both protein-protein interactions and compound-protein interactions [48, 31, 29]. Here, we use two CNN modules to extract the hidden features of peptides and proteins separately. Each CNN module consists of three convolution layers with a rectified linear unit (ReLU) function followed by a max pooling layer. The max pooling layer down-samples the output of previous filters from convolution layers to learn the features for better generalization and also reduces the output of the ReLU layer to a one-dimensional array to achieve higher learning efficiency (see Supplementary Notes S7 for more details).

#### Self-attention module

We use two self-attention modules [49] to both enhance the final binary interaction prediction performance and learn the contributions of individual residues from peptide and protein sequences to the final prediction. This attention mechanism has been commonly used in a wide variety of data analysis tasks to capture the dependencies between distant tokens in sequential inputs, in which the importance of features at each position is computed using the weighted sum over the features from all positions [49]. Here, we adopt a single-head self-attention strategy in our framework. More specifically, let ***U*** denotes the output features of the embedding layers with basic amino acid representations as input. Then the output of the single-head self-attention module is a weighted sum of the feature vectors over all residues, that is,

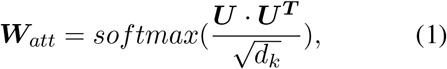

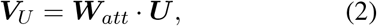

where ***W***_*att*_ represents the self-attention score matrix that implicitly indicates the contributions of features at local residues to the final prediction, ***V***_*U*_ stands for the attention matrix, and 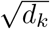 stands for a scaling factor depending on the layer size. This attention mechanism allows the model to focus on certain crucial residues from the sequences dynamically and capture the contributions of features at individual residues to the final prediction.

#### The binary interaction prediction module

The outputs of the max pooling layers from the CNN modules and the outputs of the attention modules of peptides and proteins are concatenated together and then fed into the binary interaction prediction module, which consists of three fully-connected layers. Each of the first two fully-connected layers is followed by a dropout operation to alleviate the overfitting problem. We apply a sigmoid function 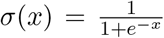 on the last layer to produce a final prediction, in which the prediction score≥ 0.5 indicates that there is an interaction between the given peptide-protein pair, and < 0.5 otherwise.

#### The peptide binding site prediction module

Given a peptide-protein pair, this module predicts which residues from the peptide sequence bind to the protein partner. The output features ***H*** of the CNN module of the peptide can be denoted by its row vectors 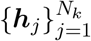, where each ***h***_*j*_ stands for the feature vector of the residue at position *j* in the peptide. We apply a single-layer neural network on ***h***_*j*_ and then normalized the output values using a sigmoid function to obtain a one-dimension value for each residue. Thus, the predicted score residue at position *j* in the peptide is

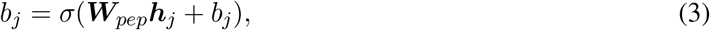

where *j* = 1, 2,…, *N*_*k*_, *N*_*k*_ represents the number of residues in the peptide sequence and *σ*(*x*) denotes the sigmoid function. Here, *b*_*j*_ ≥ 0.5 indicates that position *j* in the peptide is a binding residue, and *b*_*j*_ < 0.5 otherwise. Note that the parameters in this module are updated simultaneously with those of other modules during the training process.

### 4.5 Training

CAMP has two separate binary cross-entropy loss functions for the corresponding two classification tasks, i.e., the binary interaction prediction and the peptide binding site prediction. For the binary interaction prediction task, in a training set with *N* peptide-protein pairs, the binary cross-entropy loss is defined as

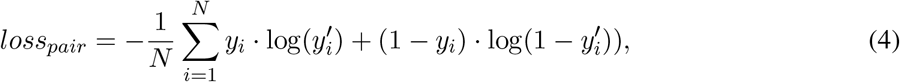

where *y*_*i*_ and 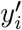 stand for the true binary label and the predicted interaction probability of a given peptide-protein pair.

For the peptide binding site prediction task, we also use a binary cross-entropy loss to measure the difference between predicted and real binding labels for individual residues in the peptide sequence. To ignore the padded zeros in our fixed-length input, we apply masks on those padded positions. More specifically, for a training set with *N* peptide-protein pairs, the masked cross-entropy loss for peptide binding site prediction is defined as

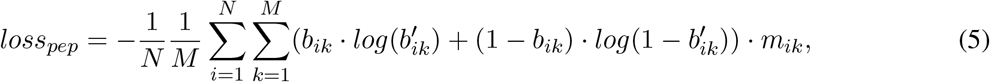

where *m*_*ik*_ stands for the mask value at position *k* in the peptide sequence of sample *i* and *M*_*i*_ = ∑*m*_*ik*_ represents the number of residues in the padded peptide sequence of sample *i* (*m*_*ik*_ is 0 if position *k* is padded with zero and 1 otherwise), and *b*_*ik*_, 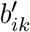 represent the true label and the predicted probability of position *k* in the *i*-th sample, respectively.

The above two losses are combined together and optimized simultaneously in a multi-objective training process, that is,

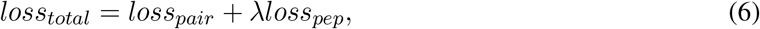

where *λ* stands for a weight parameter that balances the two losses. All parameters of CAMP are updated using the RMSProp optimizer [50]. The details about hyper-parameter tunning and selection can be found in Supplementary Notes S8. A single CAMP model can be trained within two hours on a linux server with 48 logical CPU cores and one Nvidia Geforce GTX 1080Ti GPU.

