## Supplementary files for "CAMP: a Convolutional Attention-based Neural Network for Multifaceted Peptide-protein Interaction Prediction"

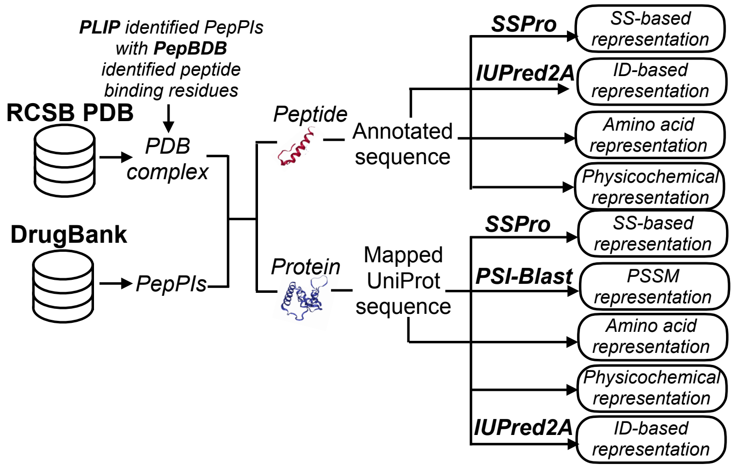


**Fig. S1.** The data curation and feature generation procedures. We first obtained all PDB complexes containing peptides as ligands from the RCSB PDB [3–5] and all peptide drugs with corresponding targets from DrugBank [6–9]. Then for the peptide-protein pairs from the PDB, we used PLIP [10] to identify non-covalent interactions. Any peptide-protein pairs involved such interactions were kept as interacting pairs. we also downloaded the corresponding labels of peptide binding residues from PepBDB [11]. Next, we mapped the protein sequences from the PDB and DrugBank to UniProt [12] to achieve unified inputs of protein sequences and then generated a series of sequence-based features for peptides and proteins including amino acid representations, secondary structure-based (SS-based) representations, physicochemical property representations, intrinsic disorder-based (ID-based) representations and PSSM representations.

### S1 Feature generation of protein and peptide sequences

**Amino acid representations of peptides and proteins** For amino acid representation, each residue is denoted by a letter from a 21-letter alphabet $\mathcal{A}$, containing a vocabulary of 21 types of amino acids (i.e., 20 canonical amino acids and a letter 'X' for any unknown or non-standard amino acid).

**Secondary structure-based representations of peptides and proteins** Here, we used SSPro [1] to predict a 3-class secondary structure type (Table S1) for each residue, which is denoted by a letter chosen from a three-letter alphabet to represent the class of the predicted secondary structure (i.e., helix, strand and the rest). Next, we define an alphabet of 63 elements to describe the combination of the predicted secondary structure type and the amino acid type of each residue. Each combination is encoded with an integer between 1 and 63.

**Physicochemical property representations of peptides and proteins** The physicochemical property representation is to encode the physicochemical features of the R group of each residue, denoted by a letter from a seven-letter alphabet based on the combination of polarity and the hydropathy index (a metric measuring the hydrophilicity and hydrophobicity of the R groups). Htis index reflects the free energy of the transfer of an amino acid side chain from a hydrophobic solvent to water [2]. The positive value indicates that the transfer is unfavorable for amino acids with nonpolar side chains and the negative value indicates that the transfer is favorable for charged or polar amino acid side chains. We therefore define an alphabet of seven elements to describe the combination of polarity and the hydropathy index of each amino acid side chain (Table S2). In particular, each combination is encoded with an integer between 1 and 7.

**PSSM representations of proteins** Given a protein sequence, we use its normalized PSSM matrix derived by PSI-blast [13, 14] (iteration = 3 and E-value=0.001). Here the parameters (the number of iterations and the threshold of E-value) have been proved to work well for generating effective feature profiles from the protein sequences [14].

**Intrinsic disorder-based representations of peptides and proteins** The intrinsic disorder scores represent the tendencies of disordered amino acid pairs to form contacts, ranging from 0 (complete order) to 1 (complete disorder). In particular, we adopt two kinds of intrinsic disorder scores, i.e., the long disorder score (considering the long disordered regions such as disordered domains) and the short disorder score (considering the short stretches of disorder such as flexible linkers and loops) [15]. We also consider the ANCHOR score (a normalized score between 0 and 1), which represents the probability of a given residue to be part of a disorder binding region, thus indicating the structure flexibility of the query residue and its neighbors [16].

**Table S1.** Representations of individual amino acid types and the secondary structure elements

| Amino Acid | | Secondary Structure | | |
|---|---|---|---|---|
| Standard Abbreviation Symbol | Unknown or Non-standard | Strand | Helix | Rest |
| Gly:G, Ala:A, ...,Glu:E | X | E | H | C |

**Table S2.** Physicochemical property representations of individual residues in protein or peptide sequences.

| Combination of Polarity and Hydropathy Index of the R Groups | | | |
|---|---|---|---|
| Polarity | Positive or Negative of Hydropathy Index | Amino-acid Members | Number of Members |
| Nonpolar | Positive | Ala, Phe, Ile, Met, Leu, Pro, Val | 7 |
| Nonpolar | Negative | Gly, Trp | 2 |
| Polar & uncharged | Positive | Cys | 1 |
| Polar & uncharged | Negative | Asn, Gln, Ser, Thr, Tyr | 5 |
| Negatively charged | Negative | Asp, Glu | 2 |
| Positively charged | Negative | Lys, His, Arg | 3 |
| Unknown | Unknown | Otherwise | NA |

### S2 Cluster-based cross-validation

In the real peptide-protein interaction prediction setting, the data redundancy problem caused by similar proteins or peptides may lead to "easy predictions", which could mislead the performance evaluation of different algorithms. To conduct an objective evaluation, we followed the same strategy as in MONN [17] and used a cluster-based strategy for cross-validation. The clustering threshold means the minimal distance between any two clusters by a single-linkage clustering algorithm [18]. More specifically, the distance between two peptides (proteins) $p_i$ and $p_j$ is defined as

$$d_{ij} = 1 - \frac{SW(p_i, p_j)}{\sqrt{(SW(p_i, p_i)SW(p_j, p_j))}}, \tag{1}$$

where $SW(\cdot, \cdot)$ stands for the Smith-Waterman alignment score (https://github.com/mengyao/Complete-Striped-Smith-Waterman-Library) between two sequences. The distance threshold values adopted in our paper were [0.3,0.4,0.5,0.6]. Here, the low threshold can ensure that the redundant sequences are separable enough while the high threshold can avoid too much imbalance between training and test data (see Table S3, S4 for more details).

In the cluster-based cross-validation procedure, we particularly chose nine-fold cross-validation for "both new setting", because under this setting, neither protein nor peptide clusters can be shared across training and test sets. In particular, we randomly split protein clusters into three folds and then divided peptide clusters into three folds inside each fold of protein clustering. Then all the peptide-protein pairs were partitioned into a $3 \times 3$ grid. We selected a single grid as the test set and removed the four grids that shared protein or peptide clusters with the chosen test set. Finally, we used the remaining four grids to train the model. In such a test setting, there was not any shared protein or peptide cluster across training set and test sets.

### S3 Ablation studies revealed that the encoded informative features and the designed network architecture of CAMP are important for peptide-protein interaction prediction

Here, we demonstrated the importance of the encoded features, the self-attention modules and the binding site prediction module by conducting a series of ablation studies. We investigated the contributions of

**Table S3.** The numbers of clusters and the max cluster sizes of proteins in our benchmark dataset under different clustering thresholds. There were in total 3412 distinct proteins in our benchmark dataset.

| Threshold | Number of Clusters | Max Cluster Size |
|---|---|---|
| 0.1 | 2822 | 24 |
| 0.2 | 2453 | 169 |
| 0.3 | 2291 | 252 |
| 0.4 | 2137 | 565 |
| 0.5 | 1981 | 566 |
| 0.6 | 1786 | 566 |
| 0.7 | 1535 | 714 |
| 0.8 | 169 | 3182 |
| 0.9 | 1 | 3412 |

**Table S4.** The numbers of clusters and the max cluster sizes of peptides under different clustering thresholds. There were in total 5399 distinct peptides in our benchmark dataset.

| Threshold | Number of Clusters | Max Cluster Size |
|---|---|---|
| 0.1 | 4270 | 76 |
| 0.2 | 3544 | 159 |
| 0.3 | 3085 | 166 |
| 0.4 | 2486 | 1209 |
| 0.5 | 1470 | 3001 |
| 0.6 | 596 | 4405 |
| 0.7 | 24 | 5309 |
| 0.8 | 1 | 5399 |
| 0.9 | 1 | 5399 |

these components by comparing the performances of CAMP on final binary interaction prediction with six alternative versions, including removing secondary structure features, physicochemical property features, PSSM profiles of the proteins, intrinsic disorder features, the self-attention modules and the peptide binding site prediction module, respectively.

**Table S5.** Performance comparison of CAMP with six alternative versions through five-fold cross-validation in a "random-split setting", in which we randomly split the benchmark dataset and used 80% of the dataset to train the model and the remaining 20% to evaluate the performance. We applied the same parameter tuning strategy as described in Supplementary Notes S8. The mean and standard deviation of each method on the benchmark dataset over five folds are shown. Also, p-values (two-tailed student t-tests of performance comparison) are reported.

| | AUC | AUPR | P-values of (AUC, AUPR) |
|---|---|---|---|
| CAMP | $\mathbf{0.8715 \pm 0.0052}$ | $\mathbf{0.6414 \pm 0.0059}$ | NA |
| -no secondary structure-based feature | $0.8597 \pm 0.0018$ | $0.6160 \pm 0.0043$ | (0.00266, 0.00011) |
| -no physicochemical feature | $0.8604 \pm 0.0065$ | $0.6263 \pm 0.0104$ | (0.02850, 0.03549) |
| -no profile-based feature | $0.8591 \pm 0.0052$ | $0.6104 \pm 0.0170$ | (0.00975, 0.02055) |
| -no intrinsic disorder-based feature | $0.8430 \pm 0.0064$ | $0.5864 \pm 0.0130$ | (0.00012, 0.00006) |
| -no attention | $0.8412 \pm 0.0227$ | $0.5810 \pm 0.0404$ | (0.03150, 0.01818) |
| -no binding site prediction | $0.8538 \pm 0.004$ | $0.6131 \pm 0.0127$ | (0.00069, 0.00373) |

Here we carried out a five-fold cross-validation procedure on our benchmark dataset and conducted two-tailed student t-tests on AUC and AUPR of each fold to quantify the difference. Table S5 showed that the secondary structure features greatly boosted the performance, and adding the evolutionary profiles of proteins as well as the intrinsic disorder scores also brought a significant improvement in pairwise interaction prediction. Although it seemed that introducing physicochemical properties of residues did not bring much increase in AUC, we observed a significant increase in AUPR. For the CAMP model without self-attention modules, both AUC and AUPR were significantly decreased. We also tested the performance of a single-objective version of CAMP, in which the peptide binding site prediction module was removed. We observed a mild decrease in the performance (i.e., 1.8% and 2.8% in terms of AUC and AUPR, respectively). This difference indicated that adding extra supervise information from peptide binding site labels can improve CAMP prediction. Furthermore, the predicted binding sites can also provide useful hints to enhance our understanding of peptide binding mechanisms. These results confirmed the effectiveness of our feature selection and important network architecture employed in CAMP.

### S4 Evaluation metrics

To evaluate the performance of CAMP on peptide binding residue prediction, we used the average AUC and average Matthews correlation coefficient (MCC) on our constructed benchmark dataset (Section 2.3 and Figure 3A and 3B) and an independent test set (Section 2.5, Figure 4C and 4D). In particular, for the peptide binding site prediction of a dataset containing $N$ peptide-protein pairs, the average AUC is defined as

$$\text{Average AUC} = \frac{1}{N}\sum_{i=1}^{N}\text{AUC}(i), \tag{2}$$

where AUC($i$) represents the area under the ROC curve of the $i$-th peptide-protein pair, which is calculated from the true binding vector $\boldsymbol{b_{pep}}$ and the predicted binding vector $\boldsymbol{b'_{pep}}$ (defined in Section 4.2 and 4.5) of the peptide.

The Matthews correlation coefficient (MCC) is defined as

$$\text{MCC} = \frac{\text{TP}\times\text{TN} - \text{FP}\times\text{FN}}{\sqrt{(\text{TP}+\text{FP})(\text{TP}+\text{FN})(\text{TN}+\text{FP})(\text{TN}+\text{FN})}}, \tag{3}$$

where TP (true positive) is the number of binding residues that are correctly predicted, TN (true negative) is the number of residues that are not involved in binding activities and are correctly predicted, FP (false positive) is the number of residues that are not involved in binding activities but are predicted as binding residues incorrectly, and FN (false negative) is the number of residues that are binding residues but are predicted incorrectly. MCC ranges from -1 to 1 and the value of zero represents a random prediction. Higher values of MCC indicate better prediction.

Then, the average MCC is defined as

$$\text{Average MCC} = \frac{1}{N}\sum_{i=1}^{N}\text{MCC}(i), \tag{4}$$

where MCC($i$) represents the Matthews correlation coefficient (MCC) of the $i$-th peptide-protein pair, which is also calculated from the true binding vector $\boldsymbol{b_{pep}}$ and the predicted binding vector $\boldsymbol{b'_{pep}}$ (defined in Section 4.2 and 4.5) of the peptide.

### S5 Performance of CAMP on predicting peptide-PBD interactions

We downloaded the data used in HSM (https://github.com/aqlaboratory/hsm), which contained 1,894,338 peptide-PBD pairs in total, covering 482 binding domains and 27,897 peptides with a positive-negative ratio of 1:37.2. Next, we conducted the same feature generation procedures as used in our original CAMP model to obtain a group of sequence-based features for both peptides and PBDs. After that, we evaluated the performance of CAMP under exactly the same eight-fold cross-validation setting as used in the HSM paper.

### S6 Supplementary details of datasets

When constructing our benchmark dataset, we excluded a specific class of antigen peptides binding to the major histocompatibility complex (MHC) molecules, since they generally possess specific binding mechanisms that may not be easily generalized to common peptide-protein interactions [19]. In addition, to maintain the high quality of the constructed dataset, we excluded those pairs that contained peptide sequences with more than 20% unknown or non-standard amino acids, or protein sequences that were longer than 5,000 amino acids as (this threshold could cover more than 99% of the protein sequences).

We set the maximum length of peptides to be 50, and those with less than 50 residues are zero-padded. Then we choose the maximum length of proteins to cover at least 80% of the intact proteins in our benchmark dataset based on the computational experiments illustrated in Table S6 and the truncating strategy from previous study [20]. Protein sequences longer than the maximum length were truncated and protein sequences shorter than the maximum length were zero-padded.

**Table S6.** The performance of CAMP on binary peptide-protein interaction prediction with different truncating lengths of the protein sequences under five-fold cross-validation. The mean and standard deviation of AUC and AUPR of each test are shown.

| Truncating Portion (%) | AUC | AUPR |
|---|---|---|
| 70 | $0.8580 \pm 0.0163$ | $0.6153 \pm 0.0249$ |
| 80 | $0.8715 \pm 0.0052$ | $0.6414 \pm 0.0059$ |
| 90 | $0.8422 \pm 0.0185$ | $0.5874 \pm 0.03165$ |
| 100 | $0.8296 \pm 0.01890$ | $0.5508 \pm 0.0392$ |

#### S6.1 RCSB PDB dataset

For each peptide-protein pair, the peptide sequence was directly obtained from the RCSB PDB with binding residues marked by PepBDB and the protein sequence was obtained by mapping to UniProt [12]. We first downloaded all complexes containing peptides as ligands from the RCSB PDB released by September 2019. Then we used the Protein Ligand Interaction Predictor (PLIP) program [10] (http://github.com/ssalentin/plip) to extract the interacting chains of peptide and protein sequences from the complex structures. Given a complex structure, PLIP recognizes seven types of non-covalent interactions, including hydrogen bonds, hydrophobic interactions, pi-stackings, pi-cations, salt bridges, water bridges and halogen bonds. A residue from the peptide and another one from the protein, with at least one non-covalent interaction was considered as an interacting pair. We then retrieved the corresponding interacting labels from PepBDB [11], a structure database of peptide-protein complexes derived from the RCSB Protein Data Bank (PDB) [3–5], which contains the peptide residues involved in hydrogen bonds and and hydrophobic contacts with the partner proteins. The peptide binding residues detected by PepBDB were then mapped to the peptide sequences (which were annotated from the RSCB PDB) using an alignment tool based on the Smith-Waterman algorithm [21] (https://github.com/mengyao/Complete-Striped-Smith-Waterman-Library). To achieve the high quality of the data, we only kept those peptide sequences with at least 80% matched residues. In total, we collected 7,233 peptide-protein pairs with 3,318 distinct protein sequences and 5,283 distinct peptide sequences, and 90.99% of the pairs had labels of peptide binding residues.

#### S6.2 DrugBank dataset

We also collected all drugs belonging to the 'peptides category' as well as their target from DrugBank [22]. For these PepPIs, peptide sequences were obtained from PubChem [23] and protein sequences were obtained by mapping to UniProt [12]. In total, we collected 196 drug-target pairs from DrugBank, including 124 and 61 protein and peptide sequences, respectively.

### S7 The convolution neural network module

Each CNN module consists of three layers including the convolution layer, the rectified linear unit (ReLU) layer and the max pooling layer. In our model, the initial feature input of the CNN module of the protein (peptide) is an $N_k \times N_d$ feature array $\boldsymbol{F}$, where $N_k$ is the length of the input protein (peptide) sequence, and $N_d$ is the dimension of features in each residue position. The convolution layer uses a sliding window of size $m$ along the array $F$ to convert $\boldsymbol{F}$ to an $[N_k] \times d$ array $\boldsymbol{H}$, where $d$ is the number of filters. Let $H_{i,k}$ represent the score of the filter $k$ for position $i$ in the array $\boldsymbol{F}$, and $W_{k,j,l}$ denote the coefficient of filter $k$ at residue position $j$ and feature $l$. Then the convolution layer computes the function

$$H_{i,k} = ReLU(\sum_{m}^{j=1} \sum_{N_d}^{l=1} W_{k,j,l} F_{i+j,l}), \quad (5)$$

where the column $X_{\cdot,k}$ is a 1-dimensional filter and $ReLU(X) = max(X, 0)$ is the activation function. Finally, the max pooling layer takes the output of the last convolution block to reduce the feature dimension.

### S8 Hyper-parameter selection

There are several hyper-parameters of CAMP, such as learning rate, the number of epochs, the number of filters and the kernel size in the convolution layers, and the size of fully connected layers. Note that due to the huge search space, it would be difficult and time-consuming to find the optimal setting of all these hyper-parameters. Therefore, we only optimize four hyper-parameters, including the number of filters in convolution layers, the size of fully connected layers, the learning rate and the coefficient of the binding site prediction loss $\lambda$ through a five-fold cross-validation procedure using a grid search approach. In particular, the search grid for the number of filters in convolution layers is [32,64,128], the grid for the size of fully connected layers is [128,256,512,1024], the grid for the learning rate is [0.0001,0.0005,0.001] and the grid for $\lambda$ is [0.01,0.1,1,10]. The hyper-parameter setting with the best AUC score over the validation set was selected. The similar searching strategy was also used for other baseline models mentioned in the Results section for a fair comparison. In particular, for DeepDTA [20], we conducted a grid search to determine the best combination of hyper-parameters, including the length of sequence window from [4,6,8,12], and we used 100 as the maximum number of epochs, which was the default value from the original paper. For PIPR [24], we conducted the same grid search strategy on the hyper-parameters as in the original paper [24], which chose the dimension of hidden states from [10,25,50] and the number of recurrent convolutional neural network (RCNN) units from [1,2,3,4]. We chose 50 as the default value for the maximum number of epoches. For NRLMF [25], we chose the optimal regulation parameters and the learning rate from [$2^{-5}$,$2^{-4}$,$2^{-3}$,$\cdots$,$2^{2}$], [$2^{-3}$,$2^{-2}$,$\cdots$,$2^{0}$], respectively.

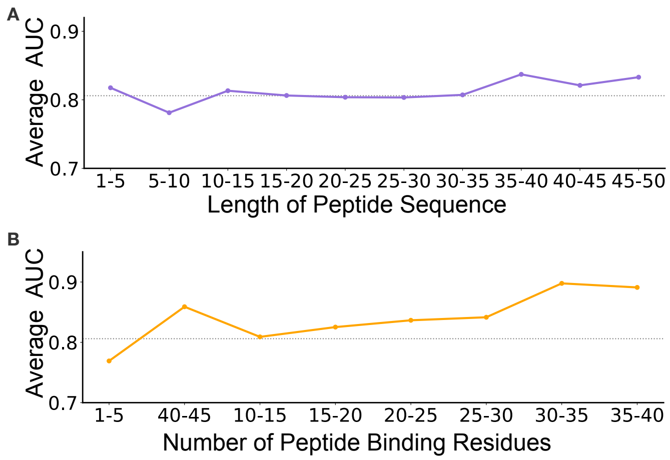


**Fig. S2.** The average AUC across various sequence lengths and numbers of peptide binding residues in the sequence. (A) shows that CAMP yields a stable performance over different lengths of peptide sequences and a slight increase of prediction scores for peptides with 35-40 amino acids, which was likely caused by a relatively larger number of samples within this length range. (B) shows that while the number of peptide binding residues increased, the average AUC displayed a slight uptrend, probably due to the extra information provided by the additional interaction information.

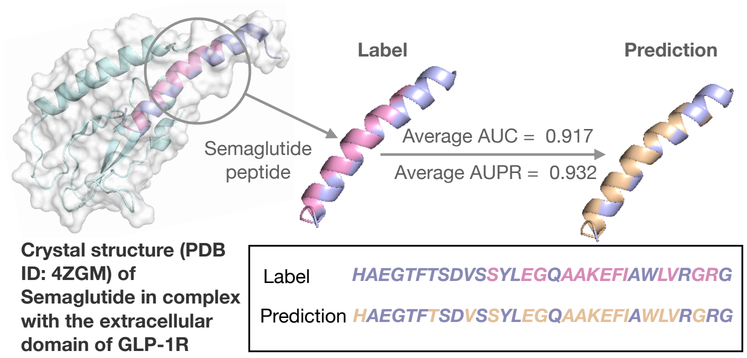


**Fig. S3.** The prediction of binding residues of Semaglutide with GLP-1R. The structure of the Semaglutide-GlP-1R complex (PDB ID: 4ZGM) was retrieved from the RCSB Protein Data Bank (PDB) [3–5] and the image was generated by PyMOL [26]. The GLP-1 receptor (UniProt ID: P43220) is colored in lightblue while Semaglutide is colored in light purple and pink. The Semaglutide peptide binds to its partner protein through 13 residues, which are colored in pink and the binding residues predicted by CAMP are colored in wheat in both sequence and structure visualization. CAMP identified 11/12 of the binding residues with five false positives.

**Table S7.** The prediction scores and ranks of GLP-1R when predicting the protein targets of Semaglutide and its analogs. ∗ denotes Semaglutide.

| PDB | Peptide Sequence | Prediction Score | Rank of GLP-1R |
|---|---|---|---|
| 3IOL | *HAEGTFTSDVSSYLEGQAAKEFIAWLVKGRG* | 0.388 | 14.47% |
| 4ZGM | *HAEGTFTSDVSSYLEGQAAKEFIAWLVRGRG*(∗) | 0.577 | 9.97% |
| 5OTT | *HXEGXFTSDLSKQMEEEAVRLFIEWLKNGGPSSGAPPPS* | 0.891 | 1.32% |
| 5OTU | *HXEGXFTSDVSSYLEGQAAKEFIAWLVKGRG* | 0.770 | 3.03% |
| 5OTV | *HCEGXFTSDVSSYLEGQAAKEFIAWLVKGRG* | 0.829 | 3.18% |
| 5OTW | *HXEGCFTSDVSSYLEGQAAKEFIAWLVKGRG* | 0.778 | 5.38% |
| 5OTX | *HCEGCFTSDVSSYLEGQAAKEFIAWLVKGRG* | 0.729 | 7.97% |
